# An evaluation of genotoxicity and 90-day repeated-dose toxicity in rats of a *Dracocephalum moldavica* powdered extract

**DOI:** 10.1101/2025.07.17.665188

**Authors:** Bean Choi, Róbert Glávits, Amy Clewell, John R. Endres, Gábor Hirka, Adél Vértesi, Erzsébet Béres, Ilona Pasics Szakonyiné

## Abstract

The safety of a powdered extract of *Dracocephalum moldavica* (Dracobelle™ Nu sd) was examined through a battery of in vitro and in vivo toxicological studies including a bacterial reverse mutation test, an in vivo mammalian micronucleus assay in male NMRI mice, and a subchronic toxicity study in Han:Wist rats. No mutagenic activity was observed in any of the tested bacterial strains with or without metabolic activation in the reverse mutation test, and no evidence of clastogenic or aneugenic activity was observed in the in vivo mouse micronucleus test. In the 90-day repeated dose oral toxicity study conducted in 80 (40 male/40 female) rats at dose levels of 0 (control), 250, 500, and 1000 mg/kg bw/day, no treatment related mortality or clinical signs of toxicity were observed, and no significant adverse effects attributable to the test item were found in any of the examined parameters in accordance with the OECD guidelines. The No Observed Adverse Effect Level (NOAEL) was determined to be 1000 mg/kg bw/day in male and female rats, the highest dose tested.

## 1. Introduction

The Moldavian dragonhead (*Dracocephalum moldavica* L., 1753) is an annual flowering plant from the Lamiaceae family, in the order of Lamiales, traditionally used by the Uygur ethnic group of Xinjiang (Maimaitiyiming et al., 2014) and in Mexico (Martinez-Vazquez et al., 2012) for various purposes. The genus consists of approximately 60 species (Zhan et al., 2024). While the plant originates from southern Siberia and central Asia, it is distributed across Egypt, China, Mongolia, and the Himalayas (Dmitruk et al., 2018; Martinez-Vazquez et al., 2012). With pale bluish-purple flowers, the genus name *Dracocephalum* is derived from the Greek words “drakon” (dragon) and “kephale” (head), referring to the shape of the corolla (Simea et al., 2018). Rich in essential oils, a strong lemon scent is characteristic of the species and is said to have a calming effect on honeybees (Dmitruk et al., 2018). In addition to essential oils, the plant contains flavonoids, terpenoids, lignans, phenylpropanoids, phenols, glycosides, polysaccharides, protein, amino acids, and minerals such as iron, copper, manganese, and strontium (Sultan et al., 2008; Zhan et al., 2024).

Due to its rich phytochemical composition, *D. moldavica* is utilized as an ingredient in food, biological pesticides, and cosmetics (Baumann et al., 2025; Pouresmaeil et al., 2022; Wandrey et al., 2021; Zhan et al., 2024). In addition to the use of *D. moldavica* leaves in snacks and bread, the seeds have been used to make snacks and tea (Dziki et al., 2019; Oniszczuk et al., 2021; Wojtowicz et al., 2017). Various extractions from *D. moldavica* have been incorporated into biodegradable packing and edible film, and a study for its potential for use as a bio-herbicide has been published (Amjadi et al., 2022; Beigomi et al., 2018; Pouresmaeil et al., 2022). Furthermore, the essential oils in *D. moldavica* are often used as ingredients in perfumes, alcoholic drinks, soaps, and detergents (Acimovic et al., 2022).

A clinical study utilizing the identical *D. moldavica* test item as used in the current work was recently published and demonstrated increased dermal thickness and improved skin hydration after oral consumption. A total of 103 subjects (aged 35–65 years) consumed 100 mg of the *D. moldavica* extract per day (or 100 mg of maltodextrin as the control) for 12 weeks. The extract was considered well-tolerated, and no adverse effects were reported during the observation period (Baumann et al., 2025).

Despite its long history of human consumption and use, *D. moldavica* has not been the subject of published comprehensive toxicological evaluations. To address this gap, we conducted a battery of toxicological studies to evaluate the safety of a powdered water *D. moldavica* extract. This paper presents the results of these studies to further contribute to its toxicological profile.

## 2. Material and Methods

### 2.1 Test Item

The test item was a yellow to brown powder of a water extract of the dry aerial parts of *Dracocephalum moldavica* (DracoBelle™ Nu sd; Mibelle Biochemistry, Buchs, Switzerland) that has been spray-dried with a maltodextrin carrier. The extract was comprised of 50% maltodextrin and 50% *D. moldavica* extract with a total flavonoid glucuronide content of ≥ 2.5 mg/g. Further characterization of the test item was performed in Baumann et al., (2025) (Baumann et al., 2025).

### 2.2 Care and use of animals

Animal studies were approved by the Institutional Animal Care and Use Committee of Toxi-Coop Zrt and conducted in accordance with The National Research Council Guide for Care and Use of Laboratory Animals (National Research Council, 2011) and the principles of the Hungarian Act 2011 CLVII (Modification of Hungarian Act 1998 XXVIII and Government Decree 40/2013). Healthy SPF male NMRI mice (Charles River Laboratories), standard animals used internationally in the mammalian micronucleus test, were acclimatized for five days. The mice weighed 37.1–41.4 g at the start of treatments and were housed 5 animals/group in Type 1 polypropylene/polycarbonate cages. The mice were group-housed to allow for social interaction and provided with deep sawdust bedding to allow digging and other normal rodent activities. Daily light exposure occurred from 6am to 6pm. The mice were kept in a controlled temperature of 22 ± 3 °C and 40–70% relative humidity.

In the 90-day study, healthy SPF Han:WIST rats (preferred and commonly used species for oral gavage toxicity studies) were acclimatized for 6 days and placed in Type III polypropylene/polycarbonate cages (2 animals/sex/cage) with certified laboratory wood bedding (SAFE 3/4-S-FASERN produced by J. Rettenmaier & Söhne GmbH+Co.KG). The body weights at the start of treatment were 183–209 g and 107–126 g for male and female rats, respectively. The cages and bedding were changed twice a week. Artificial light was maintained from 6am to 6pm (except on days of ophthalmology examinations) and temperature and relative humidity were sustained at 22 ± 3 °C and 30–70 %, respectively.

All animals were free-fed a pellet diet (ssniff SR/M-Z+H; ssniff Spezialdiäten GmbH (Experimental Animal Diets Inc.)) and had constant access to water from 250 mL bottles. Randomization of all animals to test groups was conducted using a SPSS/PC+ computer program by a trained technician. All animals were identified by unique numbers on the tails for transient identification and by ear punching for final animal numbers. Animals were selected based on a ± 20% mean body weight within each sex and a lack of clinical signs of disease or injury. Final animal numbers were given according to treatment group allocation after randomization. Selected animals were distributed by randomization according to stratification by body weight to ensure that there was no statistically significant difference among group body weight means within a sex.

### 2.3 Bacterial reverse mutation test (BRMT)

#### 2.3.1 GLP and test guidelines, test system, test material formulation, and concentrations

This GLP study followed the procedures indicated by the OECD Guidelines for Testing of Chemicals No. 471 and the ICH Guideline S2 (R1), EPA Health Effects Test Guidelines (International Conference on Harmonisation of Technical Requirements for Registration of Pharmaceuticals for Human Use (ICH) et al., 2012; OECD, 2020; U.S. Environmental Protection Agency (EPA), 1998), and established methods (Ames et al., 1975; Kier et al., 1986; Maron & Ames, 1983). The mutagenic potential of DracoBelle™ Nu sd was evaluated by measuring its ability to induce reverse mutations at selected histidine loci of bacterial strains *Salmonella typhimurium* TA98, TA100, TA1535, TA 1537, and at the tryptophan locus of *Escherichia coli* WP2 *uvrA* (All strains from Trinova Biochem GmbH; Moltox Inc.) in the presence and absence of activated rat liver S9. Based on preliminary solubility and concentration range finding tests, ultrapure water (ASTM Type 1; Toxi-Coop Zrt.) was identified as the suitable vehicle for preparing the test item solutions and concentrations of 16, 50, 160, 500, 1600, and 5000 µg/plate were prepared for the experiments.

#### 2.3.2 Experimental procedures, metabolic activation, controls, and evaluation of experimental data

A standard plate incorporation procedure was performed for the initial mutation test. Each bacterial strain was cultured in Nutrient Broth No. 2 (Oxoid) overnight and 100 μL aliquots of the culture were transferred into individual test tubes (3 tubes per control or concentration level) with 2000 μL of top agar, 100 μL of vehicle or solution of test item, 100 μL of vehicle or solution of positive controls, and 500 μL of phosphate buffer (pH 7.4; Carlo Erba; Lachner) or S9-mix. The S9-mix consisted of 500 mL of ice cold 0.2 M sodium phosphate-buffer, 100 mL rat liver homogenate (S9 fraction derived from phenobarbital and β-naphthoflavone induced rat liver (Trinova Biochem; Moltox)), and 400 mL of salt solution (β-nicotinamide adenine dinucleotide phosphate disodium salt (NADP Na; Apollo Scientific), D-glucose-6 phosphate Na (Sigma-Aldrich), magnesium chloride (MgCl_2_; Sigma Aldrich), potassium chloride (KCL; Sigma Aldrich), and ultrapure water). This solution was mixed and poured onto the surface of minimal agar plates and incubated at 37° C for 48 hours.

For the confirmatory mutation test, a pre-incubation procedure was performed. Before overlaying the test item, the bacterial culture and either S9-mix or phosphate buffer, was poured into test tubes to allow direct contact between bacteria and the test item (in its vehicle). After mixing, the test tubes were incubated for 20 min at 37° C in a shaking incubator. After the incubation period, 2 mL of molten top agar was added to the tubes and the content was mixed and poured onto minimal glucose agar plates and incubated again at 37° C for approximately 48 hours.

The tests were performed with parallel running controls: untreated, vehicle (negative), and positive control. Positive controls in the –S9 experiments, were 4-nitro-1,2-phenylenediamine (NDP; Sigma-Aldrich) for *S. typhimurium* TA98; sodium azide (SAZ; Merck KGaA) for *S. typhimurium* TA100 and TA1535; 9-aminoacridine (9AA; Merck KGaA) for *S. typhimurium* TA1537; and methyl methanesulfonate (MMS; Sigma-Aldrich) for *E. coli* WP2 *uvrA*. In the +S9 experiments, the positive control was 2-aminoanthracene (2AA; Sigma-Aldrich) for all bacterial strains. Two vehicle control groups were used depending on the solubility of the test item and the solubility of the positive control reference items. For NDP, 9AA, and 2AA, dimethyl sulfoxide (DMSO; Sigma-Aldrich) was used as the vehicle control. For the test item, SAZ, and MMS, ultrapure water was the vehicle.

Evaluation of the experimental data was conducted by counting colony numbers, comparing the data to the corresponding vehicle historical control databases, determination of cytotoxicity, and following mutagenicity criteria according to the stated guidelines.

### 2.4 In vivo mammalian micronucleus test (MMT)

#### 2.4.1 GLP and test guideline, test material formulation, administration, and dosing schedule

The in vivo MMT was GLP and OECD 474 compliant, following methods previously described (Salamone & Heddle, 1983) and the laboratory SOPs. A non-GLP preliminary oral toxicity study was conducted to determine the appropriate doses of the test item. On the day of dosing, concentrations of 50, 100, and 200 mg/mL were made with the test item and the vehicle (aqua purificata; Magilb Kft) by mixing until homogeneity was achieved and utilized within two hours.

#### 2.4.2 Experimental procedures and evaluation of data

Male NMRI mice (5 mice/group) weighing 37.1–41.4 g at the start of treatments were acclimated for five days before they were administered the test items or control (orally by gavage, twice at 24-h intervals). Cyclophosphamide (Sigma-Aldrich), the positive control, was dissolved in Aqua ad injectabilia (Magilb Kft) and administered once at a treatment volume of 10 mL/kg bw by intraperitoneal injection. Blood sampling of the negative control group occurred at 24 h after the second treatment. Clinical observations were conducted immediately after administration and subsequently at regular intervals until necropsy. Bone marrow was extracted from mice femurs to prepare for staining. Five mL of fetal bovine serum (Sigma-Aldrich) was used to flush the bone marrow which was then mixed by vortex. After concentration by centrifugation, smears were made on microscope slides and then dried at room temperature. The slides were fixed for at least five minutes in methanol (Lach:ner), stained with giemsa (Sigma-Aldrich) for 25 minutes, rinsed, dried (at least 12 hours) and coated with E-Z Mount™ (Epredia Enhancing Precision Cancer Diagnostics). The slides were coded for blind microscopic analyses. Four thousand polychromatic erythrocytes (PCEs) were scored per animal and the frequency of micronucleated cells, and the proportion of immature erythrocytes among total erythrocytes were determined.

### 2.5 90-day repeated-dose oral toxicity study in rats

#### 2.5.1 Test guideline, test system, dose administration and schedule, vehicle and test item formulation

This study was performed according to GLP, following OECD 408 Guidelines (OECD, 2018) and SOPs of the laboratory. Serum TSH level measurements were conducted in a GMP certified, non-GLP laboratory test site. Four groups of Han:WIST rats (10 rats/sex/group) were administered daily oral doses of 0, 250, 500, and 1000 mg/kg bw/day of the test item by gavage, at concentrations of 0, 25, 50, and 100 mg/mL, corresponding to a dosing volume of 10 mL/kg bw for each group. The test item and vehicle (distilled water; Magilab Kft) were administered to the appropriate animals at approximately the same time from Day 0 to Days 89/90 (males/females, respectively). Animals were not treated on the day of necropsy.

#### 2.5.2 Observations, measurements, and examinations

During the acclimation period, the eyes of all rats were examined by focal illumination and indirect ophthalmoscopy. Ten mg/mL of mydriatic eye drops (Cicloplegicedol^®^; Laboratório Edol) were administered prior to ophthalmologic examinations and the eyes were examined in subdued light by a hand ophthalmoscope. Subdued light was maintained in the rooms where animals were held for the remainder of the day. On Day 86, these eye procedures were repeated on animals of the control and high-dose groups. As there were no treatment related changes in the high-dose animals, the eyes of animals in the low– and mid-dose groups were not examined. Animals were inspected for signs of morbidity and mortality twice daily, at the beginning and end of each day.

During the treatment period clinical observations were performed cage-side after administration of the test item at approximately the same time every day. Detailed clinical observations were conducted weekly and functional observations were made prior to the termination of study. Functional observations were conducted on study Day 85 and a modified Irwin test was performed (Irwin, 1968). Body weights were recorded twice weekly during weeks 1–4 and weekly thereafter (from week 5 up to and including week 13). Food consumption was determined weekly to correspond with body weight measurements during the study. Animals were fasted overnight and blood samples were taken from the retro orbital venous plexus after Isofluran CP^®^ (Medicus Partner Kft.) administration. Clinical pathology (hematology, blood coagulation, clinical chemistry, thyroid hormone analysis), gross pathology (necropsy, organ weights) and estrous cycle examinations were conducted one day after the last treatment on Days 90/91 (male/female, respectively). Vaginal smears were prepared from each female and smears were stained with 1% aqueous methylene blue solution (VWR International LLC). Selected organs were weighed, and full histopathological examinations were performed on all animals of the control and high-dose groups (groups 1 and 4). Organs and tissues were preserved in 4% formaldehyde solution, except for testes and epididymides, which were fixed in modified Davidson solution and then stored in 4% formaldehyde solution for possible future histopathological examination. The fixed tissues were trimmed, processed, embedded in paraffin, sectioned with a microtome, placed on glass microscope slides, stained with hematoxylin and eosin, and examined by light microscopy. Additionally, histological evaluation was performed on the organs that showed macroscopic findings in the low– and mid-dose groups.

### 2.6 Statistics

Statistical analyses were conducted using SPSS PC+ software (SPSS, Inc., Version 4) and linear trends were analyzed using Microsoft Excel (version 2016). Statistical significance was set at a *p* value of < 0.05 in all studies. Barlett’s homogeneity of variance test, a one-way analysis of variance (ANOVA), Duncan’s Multiple Range test, Kolmogorov-Smirnov test, non-parametric method of Kruskal-Wallis One-Way analysis of variance, and Mann-Whitney U-tests were the statistical methods used where appropriate.

## 3. Results

### 3.1 BRMT

No biologically relevant increases in revertant colony numbers were observed in any of the five test strains following treatment with DracoBelle™ Nu sd at any concentration level, either in the presence or absence of metabolic activation. Occasional small increases in revertant colony numbers were observed throughout the experiments, however, these increases were not dose-dependent and were far below the biologically relevant thresholds for being positive. The highest mean revertant colony number increase was observed in the confirmatory mutation test in the *S. typhimurium* TA98 strain at 5000 μg/plate (-S9), with a mutation rate of 1.72, which did not meet the genotoxicological threshold for being positive. Inhibitory effects of the test item on bacterial growth were not observed in any of the examined strains in both the initial and confirmatory mutation tests and precipitation of the test item did not occur in any examined bacterial strain and concentration level (±S9).

### 3.2 MMT

No mortalities were observed in the animals and no adverse reactions to treatment were observed in either the control or test groups. DracoBelle™ Nu sd did not induce increases in the frequencies of MPCEs in male mice when compared to the concurrent negative and historical control groups. MPCEs in all treated groups were within the expected and historical control ranges. The regression analysis did not show significance, and dose-related increases were not observed. The reduction in PCE/total erythrocytes ratio was statistically lower in the 500 (compared to the historical control only), 1000, and 2000 mg/kg bw dose groups. In the 2000 mg/kg dose group, the PCE/total erythrocyte ratio showed a biologically significant 12.5% decrease, demonstrating the exposure of the test item to the bone marrow.

### 3.3 90-Day Repeated-Dose Oral Toxicity Study in Rats

#### 3.3.1 Mortality, Clinical observations, Ophthalmology, Body weights and Food Consumption

No mortalities occurred in any dose group during the 90/91 treatment period (male/female, respectively). Test item related clinical signs were not detected at any dose level during the daily clinical observations. The animals’ behavior, physiological, and neurological functions were considered to be normal at the detailed weekly clinical observations and at the functional observations in the 250, 500, and 1000 mg/kg bw/day dose groups. The body weight development was not influenced by the test item in male or female animals during the entire treatment period. There were no test item related changes in the mean daily food consumption in male or female animals. The eyes of male and female animals were normal in the control and 1000 mg/kg bw/day groups at the end of the treatment period.

#### 3.3.2 Clinical Pathology, Necropsy, Organ Weights, Histology, and Estrous cycle

No adverse effects related to the test item were observed in the hematological evaluation, blood coagulation parameters, or the clinical chemistry parameters at any examined dose at the end of the treatment period. Slight statistically significant changes in hematologic and blood coagulation parameters compared to the control were observed in the female low– and mid-dose groups, but no differences were noted in the female high-dose group (Table 1). In the low-dose group, lower mean percentage of monocytes (MONO), elevated mean percentage of basophil granulocytes (BASO) and a slightly longer mean activated partial thromboplastin time (APTT) were observed as compared to controls, and in the mid-dose group, the mean percentage of basophil granulocytes was elevated. No statistically significant hematology or blood coagulation differences compared to controls were noted in male rats of any dose group.

**Table 1.**
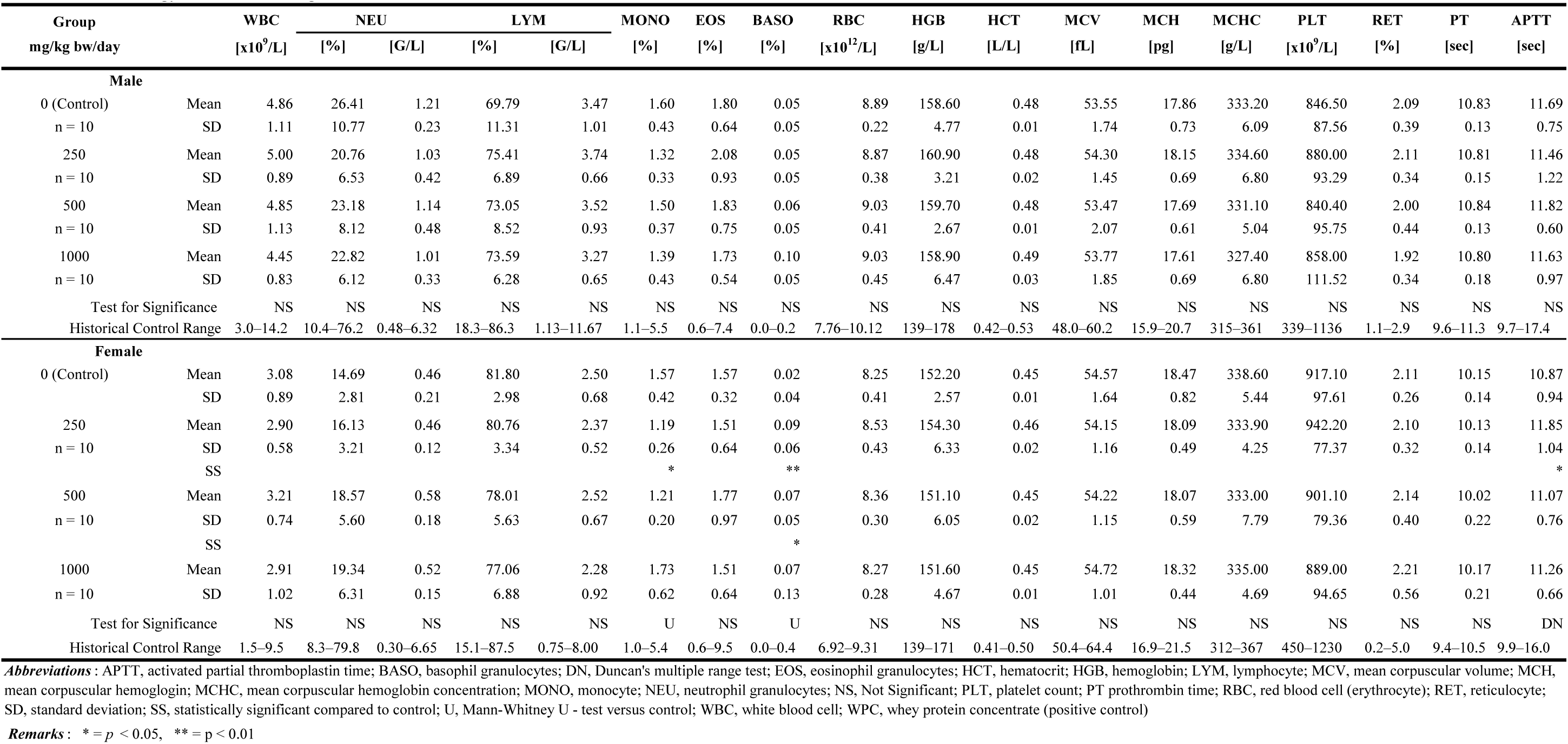
Hematology and Blood Coagulation Results.

Clinical chemistry results (as illustrated in Table 2) showed mean potassium levels (K^+^) in male animals were slightly lower than the control in the low-, mid-, and high-dose groups. In the female animals, the mean K^+^ concentration was decreased in the mid– and high-dose groups. The low-density lipoprotein concentration (LDL) and albumin:globulin ratio (A/G) in the female high-dose group were also lowered when compared to the control. The serum concentrations of the examined hormones were comparable to the control group in all examined treated dose groups (Table 3). In the high-dose male group, the FT3 concentrations exceeded the control by +22%. In the mid-dose female group, higher mean FT3 (18%) and TSH (59%) were observed, while higher mean FT3 (22%) and FT4 (23%) concentrations were detected in the high-dose groups.

**Table 2.**
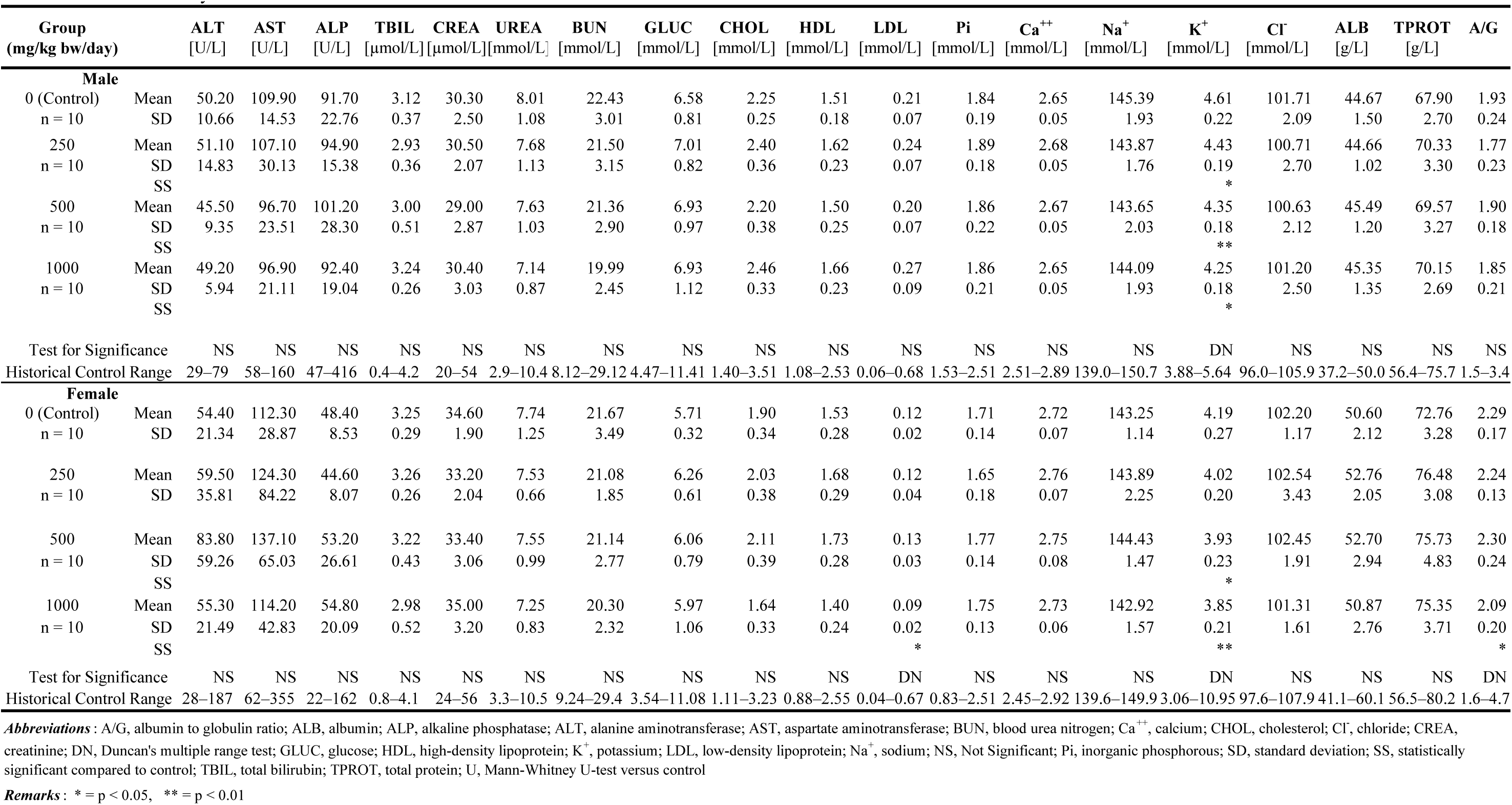
Clinical Chemistry Results.

**Table 3.**
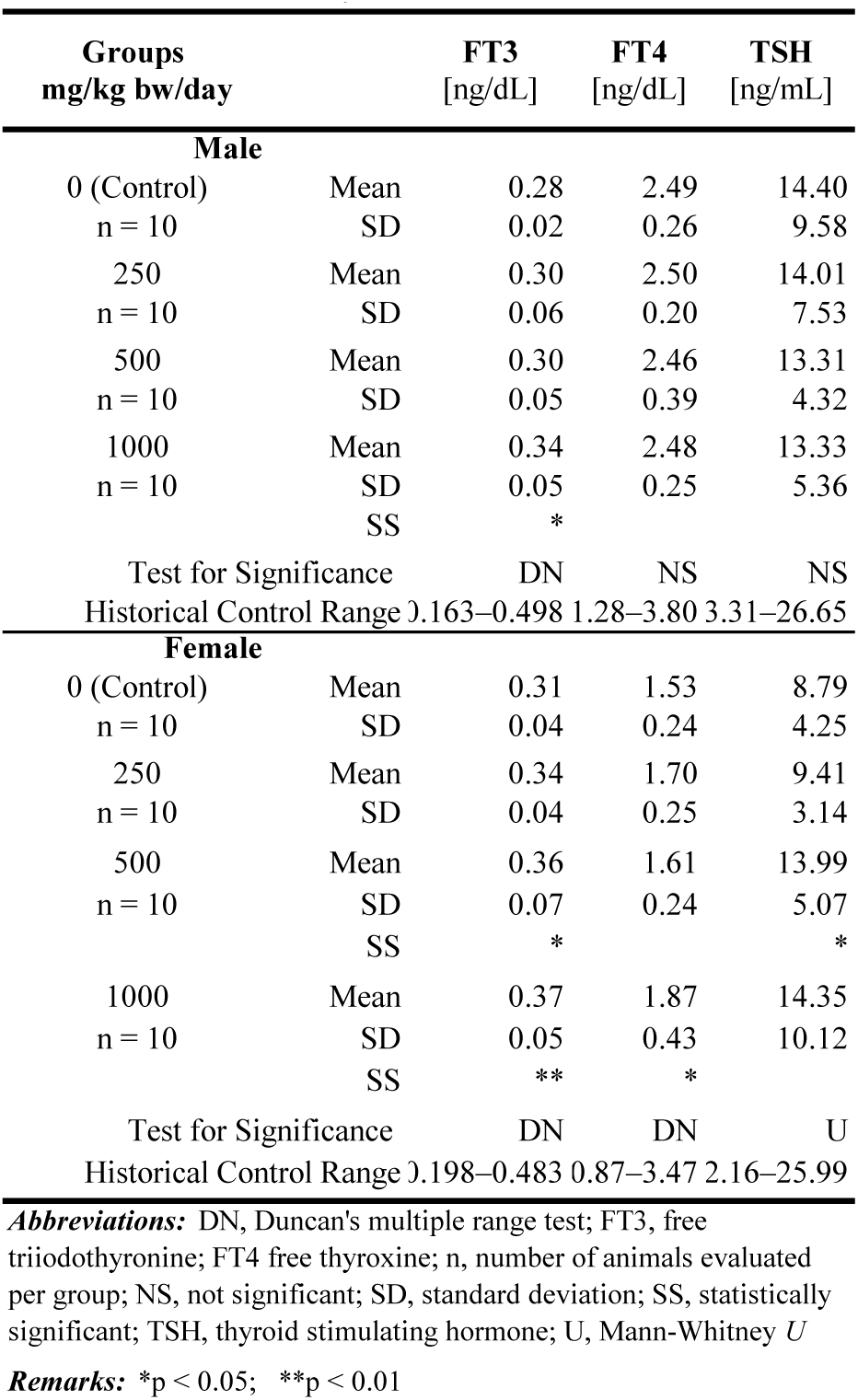
Hormone Analysis Results.

Necropsy observations did not reveal specific macroscopic lesions related to the test item in the organs or tissues at any dose level (Table 4). No test item related adverse changes in the weights of the examined organs at any dose level in male or female animals were observed. Some statistically significant differences with respect to the control were detected in the mean weights of some organs in male animals; the mean pituitary weight in the mid– and high-dose groups were decreased, the mean testes weight relative to body weight was elevated in the mid-dose group, and the mean epididymides weight relative to brain weight in the mid– and high-dose groups were elevated. Histopathological analysis did not reveal test item related lesions in the organs or tissues of animals in the high-dose group. Other histological findings consistent with macroscopic findings were judged to pose no toxicological concern (Table 5).

**Table 4.** Gross Pathology Results.

| Organs | Observations | Control | 250 | 500<br>mg/kg bw/day | 1000 |
| --- | --- | --- | --- | --- | --- |
| **Male** |  |  |  |  |  |
| Diaphragm | Hernia | 0/10 | 1/10 | 0/10 | 0/10 |
| Kidneys | No macroscopic findings | 8/10 | 7/10 | 7/10 | 8/10 |
|  | Pelvic dilation | 2/10 | 3/10 | 3/10 | 2/10 |
| Liver | No macroscopic findings | 10/10 | 9/10 | 10/10 | 10/10 |
|  | Spread into the hernia | 0/10 | 1/10 | 0/10 | 0/10 |
| **Female** |  |  |  |  |  |
| Kidneys | No macroscopic findings | 10/10 | 7/10 | 10/10 | 8/10 |
|  | Pelvic dilation | 0/10 | 3/10 | 0/10 | 2/10 |
| Uterus | No macroscopic findings | 5/10 | 8/10 | 4/10 | 4/10 |
|  | Fluid filled uterus | 5/10 | 2/10 | 6/10 | 6/10 |

**Remark :** Frequency of observations = number of animals with observations / number of animals examined.

**Table 5.** Histopathology Results.

| Organs | Observations | 0 (Control) | 250 | 500 | 1000 |
| --- | --- | --- | --- | --- | --- |
| mg/kg bw/day |  |  |  |  |  |
| **Male** |  |  |  |  |  |
| Kidneys | Pelvic dilation | 2/10 | 3/3 | 3/3 | 2/10 |
| Liver | Focal fibrosis in the Glisson's capsule | 0/10 | 1/1 | / | 0/10 |
| **Female** |  |  |  |  |  |
| Kidneys | Pelvic dilation | 0/10 | 3/3 | / | 2/10 |
| Uterus | Fluid filled uterus | 5/10 | / | / | 6/10 |

**Remarks :** Data represent number of animals with the observation per number of animals examined;
Organs without histopathology lesions in 10 of 10 control and high-dose animals or without gross lesions at necropsy not shown;
/, not examined (only gross lesions were examined in the low- and mid-dose animals)

## 4. Discussion

Minor but statistically significant findings with respect to the control were observed in the hematological, clinical chemistry, and hormone evaluations (Tables 1–3). These values were clearly within the historical control range, were without related findings, and/or were not dose-dependent and therefore were considered to be toxicologically insignificant. While a few statistically significant differences with respect to the control were detected in the mean weights of some organs in male animals, these changes were isolated, without supportive clinical pathological, macroscopic, or microscopic findings and therefore were also considered to be without toxicological relevance. Renal pelvic and uterine dilation, species-specific and frequently observed characteristics in laboratory rats, occurred in both control and test animals (Table 4) without concurrent inflammatory processes or other disease-related changes, thus were considered to be toxicologically irrelevant (Dixon et al., 1995; Johnson et al., 2013; National Toxicology Program (NTP) & DHHS, 2014). A diaphragmatic hernia that protruded into the liver (1 cm) was observed in one male from the mid-dose group and was considered to be a consistent finding in experimental rats of this strain. Histopathological findings consistent with necropsy included renal pelvic dilation, focal fibrosis in the Glisson’s capsule, and dilation of the uterine horn (Tables 4–5). The focal fibrosis of Glisson’s capsule was considered to be caused by mechanical irritation by the diaphragmatic hernia and was regarded as a developmental disorder. As stated above, renal pelvic dilation and dilation of the uterine horn are common occurrences in these rat species and considered toxicologically insignificant without other supportive findings.

To our knowledge, while much research has been conducted on the various constituents and applications of *D. moldavica* (a flowering plant native to temperate Asia and rich in phytochemicals (Acimovic et al., 2022)) no other genotoxicity or repeated-dose toxicity studies have been conducted to evaluate its safety, despite its long historical use. In the current work, we investigated the genotoxicity and subchronic toxicity of DracoBelle™ Nu sd, a powdered extract of *D. moldavica*. The test item did not elicit mutagenic activity in either the BRMT and the in vivo MMT and therefore was considered not to have shown any genotoxic activity under the conditions applied. In the subchronic toxicity study in rats, repeated exposure of the test item showed no major toxic effects and no indication of target organs. The results of these studies contribute to the safety data on *D. moldavica* available in the scientific literature.

### 4.1 Conclusions

A powdered water extract of *D. moldavica*, DracoBelle™ Nu sd, did not induce gene mutations in the genome of the strains of *S. typhimurium* TA98, TA100, TA1535, TA1537 and of *E. coli* WP2 *uvrA* in a bacterial reverse mutation assay, did not induce genotoxicity in an MMT, and did not cause adverse effects in male or female Han:WIST rats after 90/91 consecutive days of oral gavage administration of 250, 500, and 1000 mg/kg bw/day doses. The NOAEL of the subchronic study was determined to be 1000 mg/kg bw/day, the highest dose tested.

## Abbreviations

2AA: 2-aminoanthracene
9AA: 9-aminoacridine
ANOVA: analysis of variance
DMSO: dimethyl sulfoxide
DME: Dulbecco’s Modified Eagle’s
FOB: functional observation battery
GLP: good laboratory practice
ICH: International Conference on Harmonization of Technical Requirements for Registration of Pharmaceuticals for Human Use
MMS: methyl methanesulfonate
MPCE: micronucleated polychromatic erythrocyte
NOAEL: no-observed-adverse-effect-level
NPD: 4-Nitro-1,2-phenylenediamine
OECD: Organisation for Economic Co-operation and Development
SAZ: sodium azide
SOP: standard operating procedure
SPF: specific pathogen-free
TG: test guideline

## Acknowledgements

The authors thank the following individuals for their contributions to the work: participating investigators Míra Andorka, Cecília Szijártó Bándiné, Csaba Berczelly, Ibolya Bogdán, Katalin Csendes, Timea Csörge, Erika Dobos, Mónika Fekete, Zsuzsanna Frank, Petra Jáger-Gaál, Zoltán Gaál, Irén Somogyi Háriné, Ildikó Hermann, Brigitta Horváth, Istvánné Horváth, Zsolt Hummel, Ágota Jó, Bálint Zsolt Juhari, Anita Klucsik, Klára Fritz Kovácsné, Mónika Kozma, Nóra Pongrácz Kurdiné, Bence Küronya, Adrienn Laczó, Anikó Légrádi-Maurer, Marcell Madár, Máté Madár, Viktória Matina, Timea Molnár, Dániel Németh, Viktória Orovecz, Barbara Palombi, Viktória Polgár-Balogh, Anita Pélyi, Anikó Renkó, Fruzsina Ritter-Tóth, Sára Sárdi, Károly Schöll, Sebestyén Simon, David Szabó, Zsuzsanna Szabó, Mariann Lennert Szabóné, Mónika Oláh Szabóné, Edit Szám, Olga Szász, Szilvia Simai, Márta Tenk, and Erika Misku Vargáné for the performance of experimental tasks, collection of data, statistical analyses, and/or quality assurance; Jared Brodin for administrative support in preparation of the manuscript.

## Statements and Declarations

### Ethical Considerations

The animal studies were approved by the Institutional Animal Care and Use Committee of Toxi-Coop Zrt. The 90-day study was additionally conducted according to the National Research Council Guide for Care and Use of Laboratory Animals (National Research Council, 2011) and in compliance with the principles of the Hungarian Act 2011 CLVIII (modification of Hungarian Act 1998 XXVIII) and Government Decree 40/2013 regulating animal protection.

## Data Availability

The data sets generated during these studies are available on request.

## References

1. Acimovic, M., Sovljanski, O., Seregelj, V., Pezo, L., Zheljazkov, V. D., Ljujic, J., … Vujisic, L. (2022). Chemical Composition, Antioxidant, and Antimicrobial Activity of Dracocephalum moldavica L. Essential Oil and Hydrolate. Plants (Basel), 11(7). doi:10.3390/plants11070941

2. Ames, B. N., McCann, J., & Yamasaki, E. (1975). Methods for detecting carcinogens and mutagens with the Salmonella/mammalian-microsome mutagenicity test. Mutation Research, 31(6), 347–364. doi:10.1016/0165-1161(75)90046-1

3. Amjadi, S., Nouri, S., Yorghanlou, R. A., & Roufegarinejad, L. (2022). Development of hydroxypropyl methylcellulose/sodium alginate blend active film incorporated with Dracocephalum moldavica L. essential oil for food preservation. Journal of Thermoplastic Composite Materials, 35(12), 2354–2370. doi:10.1177/0892705720962153

4. Baumann, J., Bönzli, E., Wandrey, F., & Grothe, T. (2025). Moldavian Dragonhead Extract: A Natural Collagen-Booster to Target Skin Aging. OBM Geriatrics, 9(2), 17. doi:doi:10.21926/obm.geriatr.2502305

5. Beigomi, M., Mohsenzadeh, M., & Salari, A. (2018). Characterization of a novel biodegradable edible film obtained from Dracocephalum moldavica seed mucilage. International Journal of Biological Macromolecules, 108, 874–883. doi:10.1016/j.ijbiomac.2017.10.184

6. Dixon, D., Heider, K., & Elwell, M. R. (1995). Incidence of nonneoplastic lesions in historical control male and female Fischer-344 rats from 90-day toxicity studies. Toxicologic Pathology, 23(3), 338–348.

7. Dmitruk, M., Weryszko-Chmielewska, E., & Sulborska, A. (2018). Flowering and nectar secretion in two forms of the Moldavian dragonhead (Dracocephalum moldavica L.)-a plant with extraordinary apicultural potential. Journal of Apicultural Science, 62(1), 97.

8. Dziki, D., Cacak-Pietrzak, G., Gawlik-Dziki, U., Sułek, A., Kocira, S., & Biernacka, B. (2019). Effect of Moldavian dragonhead (Dracocephalum moldavica L.) leaves on the baking properties of wheat flour and quality of bread. CyTA-Journal of Food, 17(1), 536–543.

9. International Conference on Harmonisation of Technical Requirements for Registration of Pharmaceuticals for Human Use (ICH), FDA, Center for Drug Evaluation and Research (CDER), & Center for Biologics Evaluation and Research (CBER). (2012). Guidance for Industry. S2(R1) Genotoxicity Testing and Data Interpretation for Pharmaceuticals Intended for Human Use.

10. Irwin, S. (1968). Comprehensive observational assessment: Ia. A systematic, quantitative procedure for assessing the behavioral and physiologic state of the mouse. Psychopharmacologia, 13(3), 222–257.

11. Johnson, R., Spaet, R., & Potenta, D. (2013). Chapter 8. Spontaneous lesions in control animals used in toxicity studies. In Toxicologic pathology. Nonclinical safety assessment (pp. 209-254). Boca Raton: CRC Press.

12. Kier, L. D., Brusick, D. J., Auletta, A. E., Von Halle, E. S., Brown, M. M., Simmon, V. F., … Ray, V. (1986). The Salmonella typhimurium/mammalian microsomal assay. A report of the U.S. Environmental Protection Agency Gene-Tox Program. Mutation Research, 168(2), 69–240. doi:10.1016/0165-1110(86)90002-3

13. Maimaitiyiming, D., Hu, G., Aikemu, A., Hui, S. W., & Zhang, X. (2014). The treatment of Uygur medicine Dracocephalum moldavica L on chronic mountain sickness rat model. Pharmacognosy Magazine, 10(40), 477–482. doi:10.4103/0973-1296.141817

14. Maron, D. M., & Ames, B. N. (1983). Revised methods for the Salmonella mutagenicity test. Mutation Research, 113(3-4), 173–215. doi:10.1016/0165-1161(83)90010-9

15. Martinez-Vazquez, M., Estrada-Reyes, R., Martinez-Laurrabaquio, A., Lopez-Rubalcava, C., & Heinze, G. (2012). Neuropharmacological study of Dracocephalum moldavica L. (Lamiaceae) in mice: sedative effect and chemical analysis of an aqueous extract. Journal of Ethnopharmacology, 141(3), 908–917. doi:10.1016/j.jep.2012.03.028

16. National Research Council. (2011). Guide for the care and use of laboratory animals. T. N. A. Press

17. National Toxicology Program (NTP), & DHHS. (2014). Nonneoplastic Lesion Atlas. Uterus – dilation. Retrieved from https://ntp.niehs.nih.gov/nnl/female_reproductive/uterus/dilat/index.htm

18. OECD. (2018). Test No. 408: Repeated Dose 90-Day Oral Toxicity Study in Rodents, OECD Guidelines for the Testing of Chemicals, Section 4. OECD Publishing – Paris.

19. OECD. (2020). Test No. 471: Bacterial Reverse Mutation Test, OECD Guidelines for the Testing of Chemicals, Section 4. O. P.-. Paris

20. Oniszczuk, T., Kasprzak-Drozd, K., Olech, M., Wojtowicz, A., Nowak, R., Rusinek, R., … Oniszczuk, A. (2021). The Impact of Formulation on the Content of Phenolic Compounds in Snacks Enriched with Dracocephalum moldavica L. Seeds: Introduction to Receiving a New Functional Food Product. Molecules, 26(5). doi:10.3390/molecules26051245

21. Pouresmaeil, M., Sabzi-Nojadeh, M., Movafeghi, A., Aghbash, B. N., Kosari-Nasab, M., Zengin, G., & Maggi, F. (2022). Phytotoxic activity of Moldavian dragonhead (Dracocephalum moldavica L.) essential oil and its possible use as bio-herbicide. Process Biochemistry, 114, 86–92.

22. Salamone, M., & Heddle, J. (1983). Chapter 4. The bone marrow micronucleus assay: rationale for a revised protocol. In F. de Serres (Ed.), Chemical Mutagens (pp. 11–149). New York: Plenum Press.

23. Simea, Ş., Ghețe, A. B., Muresan, C., & Crișan, I. (2018). The Importance and Use of the Species Dracocepahlum Moldavica. Hop and Medicinal Plants, 26(1-2), 39–43.

24. Sultan, A., Bahang Aisa, H., & Eshbakova, K. (2008). Flavonoids from Dracocephalum moldavica. Chemistry of Natural Compounds, 44, 366–367.

25. U.S. Environmental Protection Agency (EPA). (1998). Health Effects Test Guidelines: OPPTS 870.5100 Bacterial reverse mutation test.

26. Wandrey, F., Pickel, C., Jongsma, E., Ewald, C., & Grothe, T. (2021). Evaluation of the collagen-boosting effects of a Moldavian dragonhead extract. Journal of Community Medicine and Public Health Reports, 2(7).

27. Wojtowicz, A., Oniszczuk, A., Oniszczuk, T., Kocira, S., Wojtunik, K., Mitrus, M., … Skalicka-Wozniak, K. (2017). Application of Moldavian dragonhead (Dracocephalum moldavica L.) leaves addition as a functional component of nutritionally valuable corn snacks. Journal of Food Science and Technology, 54(10), 3218–3229. doi:10.1007/s13197-017-2765-7

28. Zhan, M., Ma, M., Mo, X., Zhang, Y., Li, T., Yang, Y., & Dong, L. (2024). Dracocephalum moldavica L.: An updated comprehensive review of its botany, traditional uses, phytochemistry, pharmacology, and application aspects. Fitoterapia, 172, 105732. doi:10.1016/j.fitote.2023.105732

